# Cerebrovascular response to exercise interacts with individual genotype and amyloid-beta deposition to influence response inhibition with aging

**DOI:** 10.1101/2021.11.22.469615

**Authors:** Jacqueline A. Palmer, Carolyn S. Kaufman, Eric D. Vidoni, Robyn A. Honea, Jeffrey M. Burns, Sandra A. Billinger

**Affiliations:** Department of Physical Therapy, Rehabilitation Science, and Athletic Training, School of Health Professions, University of Kansas Medical Center, Kansas City, KS, United States of America; University of Kansas Alzheimer’s Disease Center, Fairway, KS, United States of America; Department of Molecular & Integrative Physiology, University of Kansas Medical Center, Kansas City, KS, USA

**Keywords:** cardiovascular, cognition, executive function, Stroop, Alzheimer’s disease, transcranial Doppler

## Abstract

The etiology of cognitive dysfunction associated with Alzheimer’s disease (AD) and dementia is multifactorial. Yet, mechanistic interactions among key neurobiological factors linked to AD pathology are unclear. This study tested the effect of interactions between cerebrovascular function, individual genotype, and structural brain pathology on response inhibition performance, an early and sensitive indicator of cognitive executive dysfunction with aging.

We quantified cerebrovascular response (CVR) to moderate-intensity aerobic exercise using transcranial doppler ultrasound and global amyloid-beta (Aβ) deposition using positron emission tomography in a group of cognitively normal older adults genotyped as *APOE4* carriers and noncarriers. We quantified response inhibition during a cognitive Stroop test.

Individuals with blunted CVR possessed greater Aβ deposition. There was CVR-by-carrier status-by-Aβ interaction on response inhibition. Blunted CVR was associated with impaired response inhibition specifically in carriers. Despite having greater Aβ deposition, carriers with higher CVR demonstrated better response inhibition.

Cerebrovascular interactions with individual genotype and structural brain pathology may provide a physiologically-informed target for precision-medicine approaches for early treatment and prevention of cognitive dysfunction with aging.

**Highlights:**

- Neurobiological interactions between CVR, *APOE* genotype, and Aβ are behaviorally significant.
- Blunted CVR to exercise is associated with impaired response inhibition specifically in *APOE4* carriers.
- *APOE4* carriers with more robust CVR have higher response inhibition performance, despite having greater Aβ deposition.
- Assessment of multifactorial neurobiological variables offers an early and sensitive biomarker of cognitive behavioral dysfunction with aging.

## 1. Introduction

Among a growing older adult population, the development of cognitive dysfunction is common but poorly understood. Profound individual differences in aging trajectories allow some older adults to maintain cognitive performance to the end of life while others suffer the debilitating loss of cognitive function associated with Alzheimer’s disease (AD) and related dementias (ADRD). The field of aging neurobiology increasingly recognizes cognitive decline with aging as having a mixed pathology and multifactorial etiology (Iturria-Medina et al., 2016, 2017); thus, a trans-disciplinary approach is imperative to enhance our understanding of the diverse and interacting mechanisms leading to ADRD. In the present study, we provide an initial characterization of interactions among key neurobiological factors that have been identified as significant contributors to the development of cognitive dysfunction and AD, including cerebrovascular function (Iadecola, 2004; Xie et al., 2016; Ouellette and Lacoste, 2021), structural brain pathology (Jack et al., 2010; Hampel, 2013), and individual genotype (Heffernan et al., 2016). Further, we investigate the behavioral relevance of these interactions on response inhibition performance in cognitively normal older adults.

Early detection of pathologic disease processes is imperative as a first step towards the development of effective prevention strategies and interventions for ADRD. One of the earliest behavioral manifestations of mild cognitive impairment (MCI) that can progress to AD has been identified as impaired cognitive executive function (Hutchison et al., 2010; Kirova et al., 2015). Specifically, the ability to suppress an undesired default or automatic response in the presence of interfering stimuli, or *response inhibition*, can distinguish between older adults who are cognitively normal versus MCI (Hutchison et al., 2010). Additionally, cortical activity dysfunction during response inhibition may underpin cognitive interference in motor behavior (Palmer et al., 2021), a common observation in older adults with MCI and early-stage AD (Coelho et al., 2012; Kirova et al., 2015), and older adults at high genetic risk for AD (Whitson et al., 2018). Yet the neurobiologic and physiologic mechanisms that contribute to the early-stage development of impaired response inhibition behavior remain elusive. Such knowledge could be leveraged towards the development of prevention strategies for clinical syndromes in the early stages of MCI, AD and related dementias.

Cerebrovascular pathology is commonly detected in older adults who present with behavioral deficits associated with clinical AD syndrome (Lorius et al., 2015; Wolters Frank J. et al., 2017; Sweeney et al., 2018; Bracko et al., 2021) and is linked to biomarkers of brain pathology, particularly the deposition of amyloid-beta (Zlokovic et al., 2005; Sisante et al., 2019; Solis et al., 2020). The dynamic characterization of cerebral blood flow regulation under conditions of physiologic stress enables the assessment of the cerebrovascular regulatory response to a wide variety of physiological inputs common in daily activities (Ferguson, 2014), including altered perfusion pressure, arterial blood gas, neural activity, and brain metabolism (Smith and Ainslie, 2017). Indeed, cerebrovascular regulation under such conditions of *physiologic stress* (e.g. sit-to-stand positional changes, aerobic exercise, heat stress, hypoxia) appear to play an important role in maintaining brain metabolism and function with aging (Bundo et al., 2002; Ogoh and Ainslie, 2009a, 2009b; Sato et al., 2011; Ogoh et al., 2013; Steinback and Poulin, 2016), as the effects of repeated transient disruption of blood, glucose, and oxygen supply to brain tissue accumulate over time (Tarumi and Zhang, 2018). Impaired cerebral blood flow can promote ischemic microlesions (Iadecola, 2004), and alter blood-brain barrier trafficking of amyloid-beta (Aβ) (Zlokovic et al., 2005), slowing the clearance of Aβ and promoting its accumulation in the brain.

Using transcranial Doppler ultrasound (TCD) to assess cerebral artery blood flow velocity, our group previously found that older adults demonstrate a blunted cerebrovascular response (CVR) during an acute bout of moderate-intensity aerobic exercise (Sisante et al., 2019; Alwatban et al., 2020), which may serve as an early indicator of dysfunctional cerebrovascular regulation in preclinical older adult populations (Alwatban et al., 2020). Further, CVR was associated with elevated levels of Aβ deposition, a hallmark of AD, in the brains of cognitively normal older adults (Sisante et al., 2019). Interestingly, cerebrovascular assessments performed at rest have not been able to discriminate cognitive status (Xie et al., 2016) or Aβ levels (Sisante et al., 2019) in these preclinical older adult populations, suggesting that *acute aerobic exercise assessment paradigms may enhance the detection of subtle impairments in vascular function in the early stages of disease*. Consistent with this theory, dysfunctional regulation of cerebral blood flow has been detected in older adults with MCI (Xie et al., 2016) and precedes the onset of dementia, implicating its early mechanistic role in cognitive decline with aging (Iadecola, 2004). However, despite the apparent role of CVR dysfunction in the preclinical stages of brain pathology, the interactions between CVR and other key neurobiological factors linked to early-stage cognitive impairments and AD pathology remain unknown.

Possession of *Apolipoprotein E4* (*APOE4*), the strongest known genetic risk factor for sporadic Alzheimer’s disease (Heffernan et al., 2016), is strongly linked to cerebrovascular dysfunction (Montagne et al., 2020). Cognitively normal older adults with an *APOE4* allele demonstrate a greater decline in cerebral blood flow with aging compared to noncarriers (Thambisetty et al., 2010), and an earlier blood-brain barrier breakdown that precipitates subsequent cognitive decline (Montagne et al., 2020). Recently, our group found that *APOE4* carriers with lowest resting conductance in middle cerebral artery blood flow had the greatest Aβ deposition, a relationship that was not present in noncarriers (Kaufman et al., 2021b). Yet, in the context of early-stage cognitive impairments, the behavioral relevance of APOE4 genotype in the presence of multifactorial aging pathologies among a heterogenous older adult population is poorly understood.

In this study, we aimed to 1) investigate the interactive effect of *APOE4* carrier status on the relationship between CVR to exercise and Aβ deposition and 2) test whether interactions between *APOE4* carrier status, CVR, and Aβ deposition affect cognitive performance involving response inhibition in a group of cognitively normal older adults. We hypothesized that 1) older adults with the most robust CVR to a bout of moderate-intensity aerobic exercise would have the lowest levels of Aβ deposition, with the strongest effect in *APOE4* carriers, and 2) higher CVR would attenuate the negative effect of Aβ deposition on response inhibition performance, particularly in *APOE4* carriers.

## 2. Materials and Methods

### 2.1. Participants

We selected a subset of 70 older adults from a well-characterized data registry of 125 cognitively normal older adults from the University of Kansas Alzheimer’s Disease Center (AHA 16GRNT30450008 (SB)) (Alwatban et al., 2020; Perdomo et al., 2020; Liu et al.). Our recruitment efforts within this cohort included those individuals with complete datasets for cerebrovascular function assessment, genetic *APOE* profiling, and structural neuroimaging for Aβ deposition. Inclusion criteria were (1) between 65 and 90 years old, (2) normal cognition with the absence of clinical dementia or cognitive impairment and (3) physical ability to exercise. Exclusion criteria were (1) insulin-dependent diabetes; (2) peripheral neuropathy; (3) myocardial infarction or symptoms of coronary artery disease within 2 years; (4) congestive heart failure; (5) the presence of 1 or 2 *APOE2* alleles. The experimental protocol was approved by the University of Kansas Institutional Review Board (IRB#: STUDY00001444) and all participants provided written informed consent.

### 2.2. Acute bout of aerobic exercise on a recumbent stepper

Participants underwent a bout of moderate-intensity aerobic exercise on a recumbent stepper in an experimental protocol previously described in detail (Billinger et al., 2017; Ward et al., 2018; Witte et al., 2019). Briefly, participants arrived to the laboratory between 7:30-9:00am after abstaining from caffeine for 12 hours, physical activity for 24 hours, and consuming a large meal for 2 hours. The exercise testing room was quiet, dimly lit, and temperature controlled between 22 and 24 degrees Celsius. Following 8 minutes of quiet rest seated on the recumbent stepper (NuStep T5XR), participants began an exercise familiarization bout to determine target power. Following an 8-minute seated rest recording, participants began exercising on the recumbent stepper, maintaining a step rate of 120 steps-per-minute and beginning at a resistance of 40W. The work resistance was increased until the participant reached the target heart rage range of 40-60% of age-predicted heart rate reserve and maintained this target heart rate for one continuous minute (Billinger et al., 2017; Sisante et al., 2019). The participant then maintained exercise at this moderate-intensity hear rate zone for 8 minutes.

### 2.3. Cerebro- and cardiovascular assessment and analyses

During rest and exercise bouts, we measured left cerebral blood flow velocity using transcranial Doppler ultrasound (TCD) with a 2-MHz probe (RobotoC2MD, Multigon Industries) placed over the temporal window. If the left MCA signal was not obtainable, the right MCA was used. Data were acquired through an analog to digital data acquisition unit (NI-USB-6212, National Instruments) and custom written software operating in MATLAB (The Mathworks Inc). We used a 5-lead electrocardiogram for continuous heart rate monitoring. All data were sampled at 500 Hz. Experimenters were blinded to Aβ deposition level, cognitive function, and *APOE* carrier status.

The mean MCA blood flow velocity was calculated during the 8-minute baseline rest and 8-minute moderate-intensity exercise bout. We calculated cerebrovascular response (CVR) as the difference between mean MCA blood flow velocity during exercise and mean MCA blood flow velocity at rest (Sisante et al., 2019).

### 2.4. Structural neuroimaging of amyloid-beta (Aβ) deposition

Participants underwent Florbetapir PET scans on a GE Discovery ST-16 PET/CT scanner at approximately 50 minutes after administration of intravenous florbetapir 18F-AV45 (370 MBq). Continuous acquisition of two 5-minute PET brain frames were summed and attenuation corrected (Vidoni et al., 2021). The global standardized update value ratio (SUVR) was calculated using standard procedures described previously (Liu et al.) to determine global Aβ deposition for each participant.

### 2.5. *APOE* genotyping

Whole blood was drawn and stored frozen at −80 degrees Celsius prior to genetic analyses using a Taqman single nucleotide polymorphism (SNP) allelic discrimination assay (ThermoFisher) to determine *APOE* genotype. Taqman probes were used to determine *APOE4, APOE3*, and *APOE2* alleles to the two APOE-defining SNPs, rs429358 (C_3084793_20) and rs7412 (C_904973_10) (Kaufman et al., 2021c; Vidoni et al., 2021). Individuals were classified as *APOE4* carrier in the presence of 1 or 2 *APOE4* alleles (e.g. E3/E4, E4/E4). Individuals with homozygous E3 (e.g. E3/E3) were classified as a noncarriers. Because *APOE2* is associated with reduced risk for Alzheimer’s disease, all *APOE2* carriers, whether homozygous or paired with a different *APOE* allele), were excluded from analyses in the present study (Whitson et al., 2018; Kaufman et al., 2021a).

### 2.6. Cognitive behavior assessment

#### 2.6.1. Clinical neuropsychological test battery

The Uniform Data Set (UDS) neuropsychological test battery and the Clinical Dementia Rating (CDR) scale employed by the United States Alzheimer’s Disease Research Center network (Morris, 1993; Monsell et al., 2016) were performed for all participants. All participants completed a standard in-person clinical and cognitive evaluation, during which the clinical CDR was performed by a trained clinician and the neuropsychological test battery by a trained psychometrist. Clinical and cognitive data were reviewed and finalized at a consensus diagnostic conference (Graves et al., 2015). All participants in the present analysis were rated as CDR = 0 and cognitively normal. Participant also completed a Mini-Mental State Exam (MMSE) (Folstein et al., 1975) (**Table 1)**.

**Table 1.** Participant characteristics. MMSE = Mini-Mental State Exam; Aβ = amyloid-beta; CVR = cerebrovascular response to moderate-intensity aerobic exercise. Values are depicted as mean ± SD.

|  | All<br>(n=70) | *APOE4* (+)<br>(n=20) | *APOE4* (-)<br>(n=50) |
| --- | --- | --- | --- |
| Age (years) | 71±5<br>[65-87] | 71±7<br>[65-87] | 71±5<br>[65-81] |
| Gender | F=47<br>M=23 | F=13<br>M=7 | F=34<br>M=16 |
| MMSE | 28.6±1.8<br>[22.0-30.0] | 28.6±1.4<br>[25.0-30.0] | 28.7±2.0<br>[22.0-30.0] |
| Aβ<br>Deposition | 1.09±.17<br>[0.86 to 1.6] | 1.16±.17<br>[ 0.98 to 1.6] | 1.07±.17<br>[0.86 to 1.6] |
| CVR | 5.55±5.29<br>[-13.00 to 21.88] | 5.18±4.58<br>[-3.57 to 13.78] | 5.69±5.58<br>[-13.00 to 21.88] |
| Stroop<br>ratio | 0.52±.12<br>[0.29 to 1.0] | 0.51±.12<br>[0.33 to 0.82] | 0.53±.17<br>[0.29 to 1.00] |

#### 2.6.2. Response Inhibition Performance

Participants completed a Stroop test during which they were presented with three conditions: 1) Stroop word reading, where color words were printed in black ink, 2) Stroop color naming, in which rectangular color patches were shown, and 3) Stroop interference, in which color words were printed in an incongruent ink color and participants were instructed to ignore the word and state the color of the ink. Each of the three conditions was presented individually in a 45-second trial. The raw number of correct responses was recorded within the time limit for each condition.

The difference in the time for naming the colors in which the words are printed and the same colors printed in rectangles is the measure of the interference of conflicting word stimuli upon naming colors, standardized for speed differences between people (raw number of word interference on color /raw number of control color squares read) (Stroop, J, 1935).

We calculated response inhibition performance as the Stroop ratio, where:

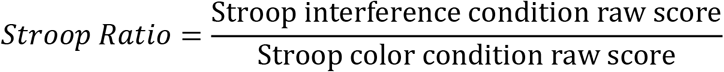

A Stroop ratio = 1.0 reflects perfect response inhibition performance and no interference of conflicting word stimuli. Here, the participant perfectly inhibited interfering word stimuli during the Stroop interference condition, with no slowing of response time relative to the control Stroop color condition. A Stroop ratio = 0.0 reflects poor response inhibition performance and complete failure to inhibit conflicting word stimuli on a color naming task. Here, the participant was unable to state any correct responses during the Stroop interference condition.

### 2.7. Statistical analyses

We tested for normality and heterogeneity of variance of all data used for analyses using Kolmogorov-Smirnov and Levene’s tests, respectively. To test the effect of *APOE4* carrier status on the relationship between CVR and Aβ, we performed two-way moderated multiple linear regression analysis after controlling for participant age. To test for interactions between *APOE4* carrier status, CVR, and Aβ deposition on response inhibition performance, we performed a three-way multiple linear regression analyses after controlling for participant age. Specifically included in the three-way model were CVR, *APOE4* carrier status, Aβ deposition, and the interaction CVR × carrier status × Aβ deposition. The relationship between Aβ deposition and response inhibition performance was compared for participants across the range of CVR to aerobic exercise. To test response inhibition performance in *APOE4* carriers and noncarriers as a function of CVR, we dichotomized participants by the group median CVR value and performed a two-way analysis of variance (ANOVA). All analyses were performed using SPSS version 25 with an a priori level of significance set to 0.05.

## 3. Results

Among the 70 total participants included in analyses (67% female, 29% *APOE4* carriers, 71±5 yo) (**Table 1**), there were no significant differences between *APOE4* carriers and noncarriers in response inhibition performance (*p*=.301), age (*p*=.94), gender distribution (*p*=.723), MMSE (*p*=.916), or CVR to aerobic exercise (*p*=.716) in carriers and noncarriers. *APOE4* carriers had greater levels of Aβ deposition compared to noncarriers (*p*=.02).

### 3.1. Relationship between cerebrovascular function and amyloid beta and effect of *APOE4*

After controlling for age, the two-way multiple regression model significantly predicted Aβ deposition, F_4,69_=4.97, p=0.001, R^2^=0.23, adjusted R^2^=0.19. Regression coefficients and standard errors are detailed in **Table 2**. A higher CVR predicted lower Aβ deposition across all participants (*p*=0.013) (**Figure 1**). Though *APOE4* carriers showed a stronger relationship between CVR and Aβ deposition (r=0.50) than noncarriers (r=0.38), the difference of this relationship was not significant (t=−0.65, *p*=.517). There was a main effect of *APOE* carrier status on Aβ deposition in the model, with *APOE4* carriers having greater Aβ deposition than noncarriers (*p*=.042) (**Table 2)**.

**Figure 1.**
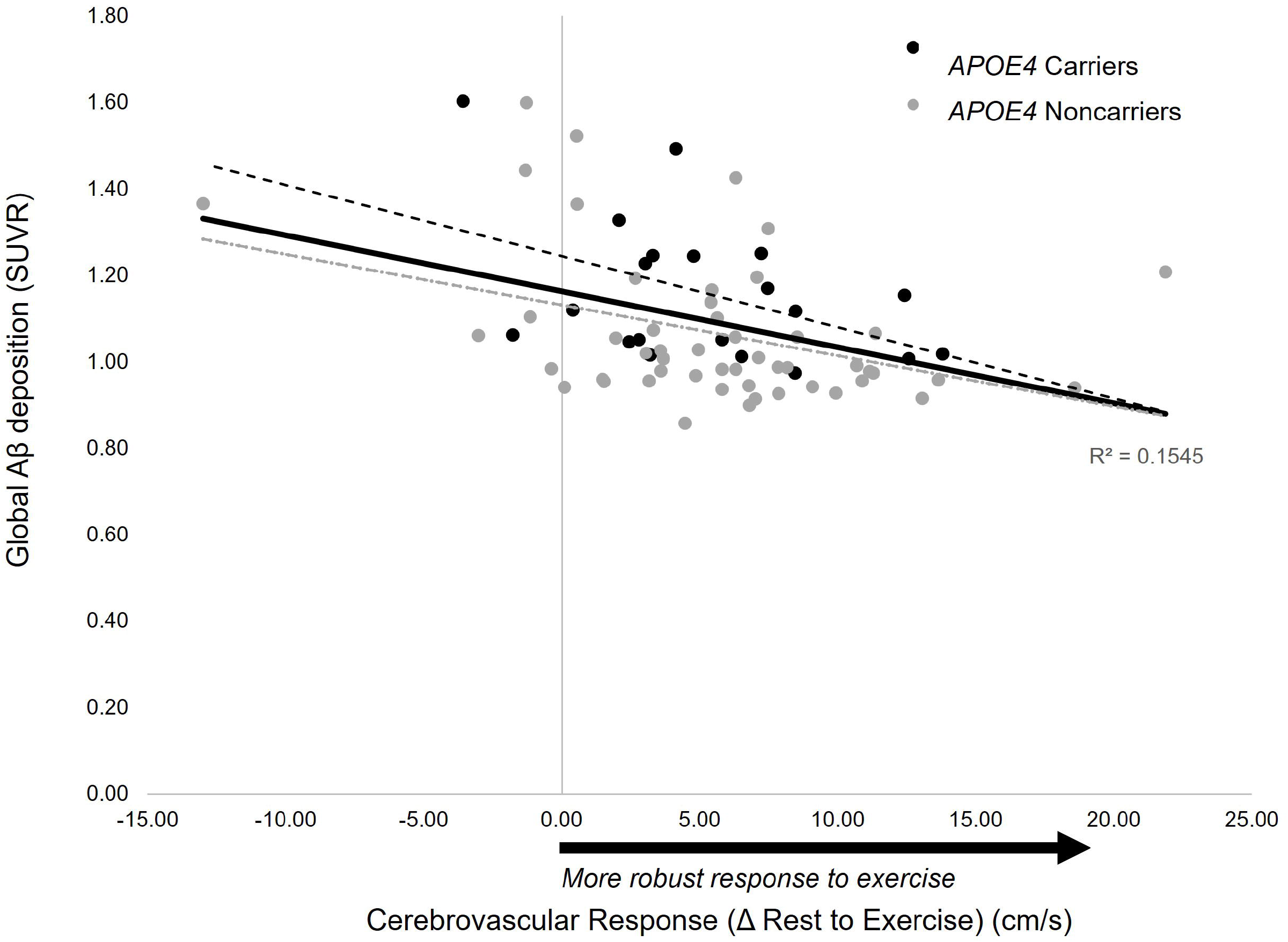
Relationship between cerebrovascular response to moderate intensity exercise and global amyloid-beta (Aβ) burden. A higher CVR predicted lower Aβ deposition across all participants (*p*=.013).

**Table 2.** Two-way moderated multiple regression analysis testing the effect of *APOE4* carrier status on the relationship between CVR and Aβ deposition after controlling for age, N = 70; Regression model predicting Aβ deposition, *p*= .0008.

| Variable | *B* | *SE<sub>B</sub>* | $\beta$ | p-value |
| --- | --- | --- | --- | --- |
| Intercept | 0.718 | -0.257 |  | .007* |
| Age | 0.005 | 0.004 | 0.152 | .176 |
| CVR | -0.058 | 0.023 | -0.319 | .013* |
| *APOE4* carrier status (+) | 0.087 | 0.042 | 0.225 | .042* |
| Interaction *APOE4* carrier status (+) × CVR | -0.032 | 0.049 | -0.080 | .517 |

Aβ = amyloid-beta; CVR = cerebrovascular response to moderate-intensity aerobic exercise; $SE_B$ = standard error of the coefficient; *B* = *unstandardized regression coefficient*; $\beta$ = standardized coefficient; \* $p<.05$

### 3.2. Interactions between cerebrovascular function, amyloid-beta, and APOE genotype as predictors of response inhibition behavior

The multiple regression model significantly predicted response inhibition performance, *F*_5,69_=3.55, *p*=.007, *R*^2^=0.22, adjusted R^2^=0.16. Regression coefficients and standard errors are detailed in **Table 3**. There was a significant three-way interaction between CVR, *APOE* carrier status, and Aβ deposition (t=2.79, *p*=.007), indicating differences in the predictive value of Aβ deposition and CVR on response inhibition performance for *APOE4* carriers and noncarriers. Specifically, *APOE4* carriers with higher CVR demonstrated better response inhibition performance (*p*<.001), while noncarriers showed no significant effect of CVR (*p*=. 112) (**Figure 2**). The model showed that *APOE4* carrier status moderated the interactive effect of CVR and Aβ deposition on response inhibition, as illustrated in **Figure 3A & B**. *APOE4* carriers who achieved a high CVR showed a positive relationship between Aβ deposition and response inhibition, where carriers with a robust CVR to an acute bout of aerobic exercise performed better on the response inhibition task despite having high levels of Aβ deposition. In contrast, *APOE4* carriers who had lower CVR showed no relationship **(Figure 3A)**. Noncarriers showed less effect of CVR on the relationship between AB deposition and response inhibition, where older adult noncarrier cognitive performance was independent of their CVR to the exercise bout **(Figure 3B)**.

**Figure 2.**
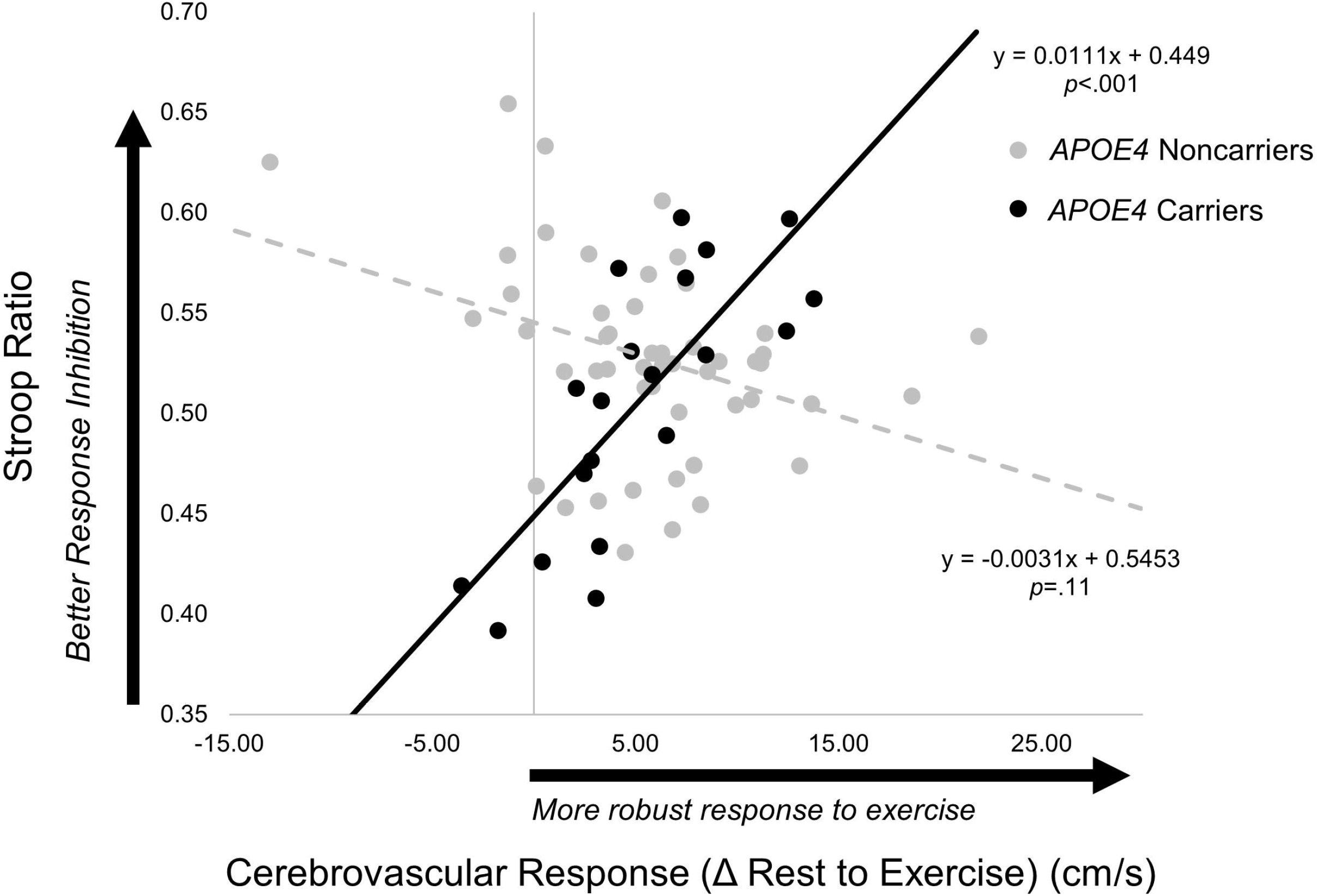
Association between cerebrovascular response to moderate-intensity aerobic exercise and response inhibition performance predicted from the regression model in *APOE4* carriers (n=20) (solid line) and noncarriers (n=50) (broken line). In *APOE4* carriers, greater cerebrovascular response to exercise was associated with better performance on the Stroop response inhibition task (*p*<.001). In noncarriers, no relationship was observed (*p*=.112). Results for this regression model are detailed in Table 3.

**Figure 3.**
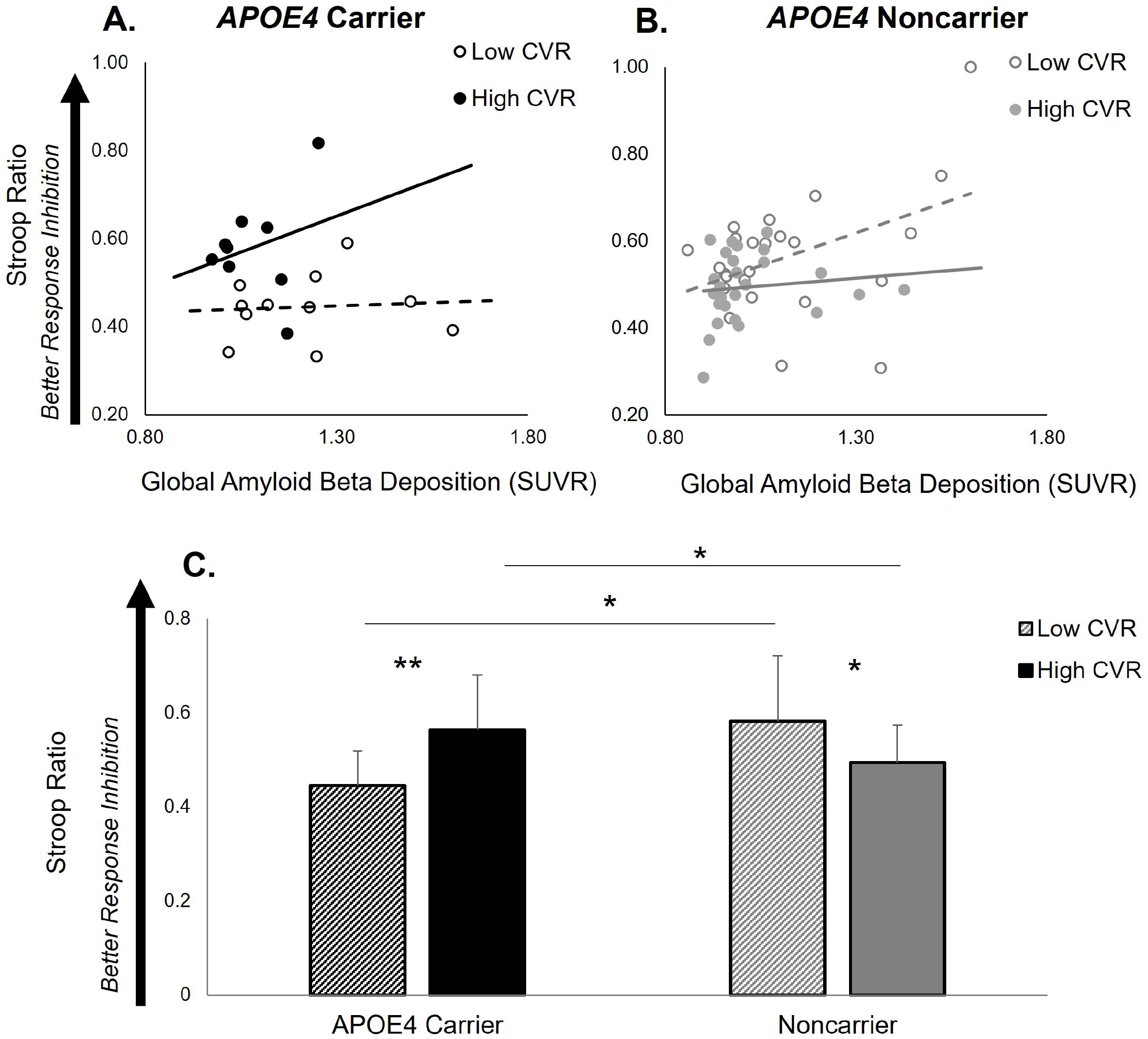
Association between Aβ deposition and response inhibition performance illustrated as a function of cerebrovascular response (CVR) to moderate-intensity aerobic exercise in *APOE4* carriers (black color) (n=20) (A) and noncarriers (grey color) (n=50) (B). *APOE4* carriers with high CVR (≥5.7cm/s) (solid line) showed a positive relationship between Aβ deposition and response inhibition performance, while APOE4 carriers with low CVR (< 5.7cm/s) (broken line) showed no positive effect. CVR also showed a lower effect on response inhibition in noncarriers. Results for this regression model are detailed in Table 3. (C) Response inhibition performance in *APOE4* carriers and noncarriers with low and high CVR. There was a significant *APOE4* carrier group × CVR interaction. *APOE4* carriers with high CVR had higher response inhibition performance compared to *APOE4* carriers with low CVR (*p*=.005) and noncarriers (*p*=.017). Among individuals with low CVR, APOE4 carriers had lower response inhibition performance compared to noncarriers (*p*=.013). Within the noncarrier group, noncarriers with low CVR had higher response inhibition performance compared to those with high CVR (*p*=.034). *p<.05, **p<.01. CVR is illustrated as a median (5.7cm/s) split of the group, where High CVR ≥ 5.7cm/s; Low CVR < 5.7cm/s.

**Table 3.** Three-way multiple regression analysis results to predict response inhibition performance after controlling for age, N = 70; Regression Model, *p*=.007.

| Variable | $B$ | $SE_B$ | $\beta$ | p-value |
| --- | --- | --- | --- | --- |
| Intercept | 0.680 | 0.190 |  | .001* |
| Age | -0.006 | 0.002 | -0.258 | .029* |
| CVR | -0.002 | 0.003 | -0.079 | .557 |
| *APOE4* carrier status (+) | -0.130 | 0.043 | -0.511 | .004* |
| Aβ deposition | 0.236 | 0.084 | 0.355 | .007* |
| Interaction *APOE4* carrier status (+) * Aβ * CVR | 0.015 | 0.005 | 0.485 | .007* |

Aβ = amyloid-beta; CVR = cerebrovascular response to moderate-intensity aerobic exercise; $SE_B$ = standard error of the coefficient; $B$ = *unstandardized regression coefficient*; $\beta$ = standardized coefficient; \* $p<.05$

When the group was dichotomized by the group median CVR value (5.7 cm/s), there was a significant *APOE4* carrier group × CVR interaction ((F_3,69_)=12.84, *p*=.001) on response inhibition performance **(Figure 3C)**. Post-hoc analyses revealed that *APOE4* carriers with a high CVR ( ≥ 5.7cm/s) demonstrated higher response inhibition performance compared to *APOE4* carriers with low CVR (< 5.7cm/s) (*t*= −3.19, p=.005) and noncarriers with high CVR (*t*=−2.51, *p*=.017). Among participants with low CVR, *APOE4* carriers had poorer response inhibition performance compared to noncarriers (*t*=2.64, *p*=.013). Within the noncarrier group, noncarriers with low CVR had higher response inhibition performance compared to those with high CVR (*t*=2.18, *p*=.034) **(Figure 3C)**.

## 4. Discussion

Findings of the present study shed light on the complex neurobiological interactions of factors known to influence the development of cognitive dysfunction with aging. Our results provide novel evidence for the behavioral significance of interactions between cerebrovascular function, individual genotype, and structural brain pathology. Despite having higher Aβ deposition, we found that *APOE4* carriers who had a more robust CVR to exercise had higher response inhibition performance, an effect that was not present in noncarriers. These findings imply higher cerebrovascular function may serve a neuroprotective role for the preservation of cognitive executive function specifically in *APOE4* carriers, possibly through increased resiliency to Aβ deposition with aging. Building upon previous research, our findings support an individualized framework gleaned from a multifactorial approach for detection of subtle impairments in cognitive behavior prior to the onset of clinical syndrome associated with AD and related dementias. Together, these results may be informative for targeted precision-medicine approaches for early intervention and prevention of age-related declines in cognitive executive function.

### 4.1. Cerebrovascular response (CVR) to exercise is linked to amyloid beta (Aβ) deposition in the absence of clinical cognitive dysfunction

Our findings support the assessment of CVR to acute aerobic exercise as a useful functional biomarker of structural brain pathology in the absence of clinically detectible impairments in cognitive function. In the present study, we found that cognitively normal older adults with blunted CVR to aerobic exercise demonstrated greater Aβ deposition (**Figure 1**), a finding consistent with previous work from our laboratory (Sisante et al., 2019) across a larger cohort of older adults. This finding also supports previous research implicating cerebrovascular assessments performed under conditions of physiologic stress may be sensitive enough to detect the first signs of dysfunction in the regulation of cerebral blood flow that may be salient to subtle changes in cognitive function in preclinical older adult populations. Although a more robust CVR demonstrated a stronger association with lower Aβ deposition in *APOE4* carriers compared to noncarriers (i.e. steeper negative slope in *APOE4* carriers (**Figure 1)**), the influence of *APOE4* carrier status on the negative association between CVR and Aβ deposition was not significant, in contrast to our a priori hypotheses (**Table 2**). This may be explained by the exclusion of older adults with MCI and a history of cardiovascular disease, which effectively narrowed the range of CVR and Aβ deposition in our participant cohort. In future studies that expand cognitive and cerebrovascular inclusion criteria, differences in this relationship between *APOE4* carriers and noncarriers may emerge, in which the effect of negative association between CVR and Aβ deposition may be strongest in *APOE4* carriers.

Our findings implicate common neurobiological mechanisms that contribute to cerebrovascular dysfunction and structural brain pathology in the early stages of brain aging. Our results demonstrate a negative relationship between CVR to exercise and Aβ deposition that is consistent with a mechanistic link between CV dysfunction and deposition of Aβ in the aging brain. However, the causal nature of this relationship remains to be tested. For example, the presence of Aβ may blunt the CVR to exercise through increased resistance of blood flow and perfusion to cortical tissues as blood flow velocity increases during aerobic exercise (Niwa et al., 2002; Iadecola, 2004). Conversely, poor cerebrovascular function may alter blood-brain barrier trafficking of Aβ, slowing the clearance of Aβ and promoting its accumulation in the brain (Zlokovic et al., 2005). The later may be supported by evidence that habitual aerobic exercise appears to have no effect on Aβ deposition in older adults (Vidoni et al., 2021), yet can improve cerebrovascular health (Thomas et al., 2013; Whitaker et al., 2020) and slow or, in some cases, reverse the development of cognitive impairment in older adults (Zhang et al., 2020). These observations may explain the neuroprotective effect of aerobic exercise against AD and related dementias (Tarumi and Zhang, 2018). Future research aimed at modulating each of these variables (e.g. aerobic exercise interventions that improve CVR) will provide insight into the directionality of the relationship between cerebrovascular function and structural brain pathology.

### 4.2. Behavioral significance of interactions between individual genotype, cerebrovascular function, and brain structure

Importantly, our findings provide evidence for the *behavioral significance* of interactions between individual genotype, cerebrovascular function, and brain structure. We found that cerebrovascular function interacts with individual *APOE* genotype and Aβ deposition to influence an individual older adult’s ability to inhibit a default and undesired response (**Figure 2** and **Figure 3A&B**). Our results are consistent with accumulating evidence within the field of aging neurobiology and the increasing recognition of the mixed pathology and multifactorial etiology that contributes to cognitive decline with aging (Iturria-Medina et al., 2016, 2017). Clinically, it is well known that an individual who has structural brain biomarker evidence of AD pathology (e.g. high levels of Aβ deposition) may paradoxically present with the absence of any behavioral indicator associated with AD clinical syndrome. Notably, vascular pathology and individual genotype (e.g. possession of the *APOE4* allele) are frequently detected in the typical clinical presentation of AD (Iadecola, 2004; Xie et al., 2016; Ouellette and Lacoste, 2021), and can significantly increase an individual’s risk for developing dementias (Heffernan et al., 2016). Building upon our current knowledge of each of these factors individually, our present results provide unique insight into interactions between these key neurobiological factors that can be gleaned from an interdisciplinary and multifactorial assessment of AD and related dementia risk factors. This rich multifactorial approach offers a powerful characterization of factors potentially influencing the early development of cognitive behavioral impairment that could have clinically meaningful implications for intervention, treatment, and prevention of AD and related dementia in preclinical older adult populations.

### 4.3. Role of cerebrovascular health as a function of individual genotype in the preservation of brain behavioral function with aging

Our results suggest cerebrovascular function plays a key role in the early behavioral manifestations of cognitive executive dysfunction in older adults who carry the *APOE4* allele. The three-way interaction model found that older adult *APOE4* carriers with blunted CVR to exercise demonstrated poorer response inhibition performance, while CVR did not predict behavioral performance in noncarriers (**Figure 2**). The characterization of these interactions builds upon previous findings from our laboratory, in which we demonstrated measures poor cardiovascular health negatively influenced the CVR to exercise preferentially in *APOE4* carriers (Kaufman et al., 2021c). Our findings are also consistent with previous evidence demonstrating a stronger effect of cardiovascular health on cognitive performance metrics in *APOE4* carriers (Zade et al., 2010; Caselli et al., 2011; Shaaban et al., 2019). Specifically within a behavioral context, a key novel finding of the present study identifies the unique relationship between cerebrovascular regulation and response inhibition performance in *APOE4* carriers compared to noncarriers. The positive relationship between CVR and response inhibition performance supports that *higher cerebrovascular regulatory function may serve a unique neuroprotective function in early brain aging processes affecting cognition in APOE4 carriers*, while playing less of a role in noncarriers. Interestingly, Kaufman et al. (2021a) revealed that cerebrovascular dysfunction in *APOE4* carriers was *differentially modifiable* through an aerobic exercise intervention compared to noncarriers, with older adult *APOE4* carriers showing greater improvements in cerebral blood flow following a year-long exercise intervention (Kaufman et al., 2021a). Together, these findings identify CVR to exercise as a useful early assessment tool in preclinical older adult populations and may provide a physiologically-informed target for precision-medicine approaches aimed at preventing age-related declines in cognitive executive function.

### 4.4. Genetic mechanisms influencing interactions between cerebrovascular function and structural brain pathology in the context of cognitive behavior

Possession of an *APOE4* allele may reveal unique genetic mechanisms that influence interactions between cerebrovascular regulation and structural brain pathology to influence cognitive behavior in older adults. Specifically, we found differential effects of CVR on the relationship between Aβ and response inhibition in *APOE4* carriers and noncarriers (**Figure 3A&B**). In *APOE4* carriers, individuals with a higher CVR and greater Aβ deposition had higher response inhibition performance, an effect not observed in *APOE4* carriers with low CVR or noncarriers (**Figure 3A&B**). Further, *APOE4* carriers with high CVR outperformed both carriers with low CVR and noncarriers on the response inhibition task (**Figure 3C**). In the face of age-related neuropathology, some individuals appear to utilize a neurologic “reserve” that enables behavioral compensation and attenuates cognitive decline (Stern et al., 2019). Findings in the present study provide preliminary evidence that the neuroprotective effect of high CVR in APOE4 carriers may act mechanistically through increased resilience to age-related structural pathology reflected in Aβ deposition, potentially supporting a functional neurologic “reserve” in the aging brains of these individuals. We also found that, among participants with low CVR, *APOE4* carriers had poorer response inhibition compared to noncarriers (**Figure 3C**). This suggests that cerebrovascular health not only serves a specialized protective effect to *maintain* cognitive function, but may additionally *attenuate the rate of cognitive decline* that is accelerated in *APOE4* carriers (Caselli et al., 2004, 2007). It is conceivable that such differences in the resiliency effect of cerebrovascular function between *APOE4* carriers and noncarriers may overlap with the neurobiological mechanisms that predispose *APOE4* carriers to a higher genetic risk for the development of AD in the later stages of aging (Farrer et al., 1997; Heffernan et al., 2016). Further, our findings that CVR has a greater effect on cognitive function in *APOE4* carriers compared to noncarriers supports the notion of different etiologies and pathologic disease processes leading to the development of AD for each *APOE4* carriers and noncarriers (Emrani et al., 2020), particularly that cerebrovascular dysfunction may play a preferential role in dementia pathogenesis for carriers (Høgh et al., 2001; Montagne et al., 2020). These preliminary findings motivate future research in a larger cohort of heterogeneous older adults that may additionally include *APOE2* carriers, for whom the effect of resiliency to aging processes has been most consistently observed and reported (Farrer et al., 1997; James et al., 2017).

While high cerebrovascular function demonstrated a neuroprotective effect for *APOE4* carriers, an interesting pattern in our data suggests that noncarriers may possess greater resiliency to potential negative effects of low CVR and high Aβ deposition. While *APOE4* carriers with high CVR had higher response inhibition performance, noncarriers with *low* CVR tended to have higher response inhibition performance than noncarriers with high CVR, though the relationship of this effect did not meet our a priori adopted level of significance **(Figure 2**) and showed a lower magnitude of effect compared to the contrasting effect observed in *APOE4* carriers (**Figure 3C**). Previous studies found that, compared to cognitively normal *APOE4* carriers, noncarriers showed differences in cortical neural function (Leuthold et al., 2013) and had greater adaptive processes to repeated trauma (James et al., 2017). Although the mechanisms are not yet fully understood, these interactive functional and structural processes appear to increase neuroprotection and promote brain resilience to specific disease processes in noncarriers, including the development of MCI (Pa et al., 2009), posttraumatic stress disorder (Peterson et al., 2015), and recovery from traumatic brain injury (Zhou et al., 2008). A greater ability to repair and protect against neuronal damage offered by the noncarrier-specific genotypic interaction (Mahley and Rall, 2000) may similarly increase noncarriers’ resilience to the chronic neurotoxic effects poor cerebrovascular function on both the accumulation of Aβ in the brain (**Figure 1**) and cognitive performance (**Figure 2** and **Figure 3**) compared to *APOE4* carriers in the present study. Notably, our exclusion of older adults with MCI would have biased the selection of only individuals with such genotypic resiliency (i.e. excluding those with low CVR and high Aβ without this protective adaptive characteristic). The present participant selection criteria may explain poorer response inhibition performance in noncarriers with low Aβ and high CVR (**Figure 3B & C**). Future research in animal models may elucidate our mechanistic understanding of risk and resilience associated with individual *APOE* genotype and further identify epigenetic pathways underpinning individual-specific physiologic stress responses for neuronal repair and protection in the context of cerebrovascular health and cognitive dysfunction with aging.

### 4.5 Cerebrovascular dysfunction affects neural networks involved in response inhibition in the early stages of cognitive dysfunction with aging

Our hypothesis-driven analyses in the present study focused on the neurobiological factors influencing a highly specific aspect of cognition involving inhibition of a default and undesired response(Stroop, J, 1935; Hutchison et al., 2010). Yet, previous research supports the specificity of cerebrovascular function and individual *APOE* genotype (Whitson et al., 2018) effects on response inhibition behavior that engages inhibitory neural networks within the prefrontal cortex (PFC) (Wessel and Aron, 2017; Wessel et al., 2019). In neurologically-intact young adults, differences in acute cerebrovascular changes within the frontal cortical regions could be dissociated during a challenging Stroop task; here, participants with a greater frontal cortical hemodynamic response demonstrated higher response inhibition performance (Gratton et al., 2020). In aging populations, previous studies have demonstrated n that cortical inhibitory function is one of the earliest neural mechanisms affected by aging processes in cognitively normal older adults (Nielson et al., 2002; Heise et al., 2013; Levin et al., 2014; Rossiter et al., 2014; Legon et al., 2016), which may explain the presence of impaired response inhibition in older adults with absent clinical cognitive syndrome in the present study. Interestingly, PFC brain regions may preferentially benefit from the therapeutic effects of aerobic exercise interventions on cognition (Duchesne et al., 2015; Levin and Netz, 2015), supporting the robust interactive effect of cerebrovascular function on response inhibition performance in the present study. Taken together, these findings may inform the development of individualized approaches for therapeutic noninvasive brain stimulation (e.g. neuromodulation of PFC brain regions in preclinical *APOE4* carriers) for the treatment and prevention of cognitive dysfunction with aging (Hsu et al., 2015).

## 5. Conclusions

Interactions between cerebrovascular function, individual genotype, and structural brain pathology may offer a useful and sensitive biomarker for early preclinical behavioral manifestations of cognitive impairment in older adults. Additionally, cerebrovascular response to aerobic exercise may provide a physiologically-informed target for precision-medicine approaches aimed at attenuating negative effects of structural brain pathology and preventing age-related declines in cognitive executive function. Future research may reveal that individuals with *APOE4* carrier genotype show the highest therapeutic benefit for such intervention approaches.

## Abbreviations

CVR: cerebrovascular response
Aβ: amyloid-beta
APOE4: Apolipoprotein E allele 4
AD: Alzheimer’s Disease
MCI: mild cognitive impairment
ANOVA: analysis of variance
SD: standard deviation

## Author Contributions

JP conceived of the presented research question and designed the analysis for this study. JB and SB provided oversight to the data collected and supervised all data analyses. CK contributed to recruitment and data collection. JP constructed the figures and the first draft of this manuscript. EV and SB contributed the analysis tools for cerebrovascular data. CK and RH contributed analysis tools, oversight and interpretation for the individual genotyping. SB supervised the findings of this work. All authors discussed the results and contributed to the final manuscript.

## 1 Data Availability Statement

The datasets generated and analyzed supporting the conclusions of this article will be made available by the authors upon request, without undue reservation.

